# Molecular confirmation of *Anopheles stephensi* in the Al Hudaydah Governorate, Yemen, 2021-2022

**DOI:** 10.1101/2024.02.22.577782

**Authors:** Methaq Assada, Mohammed Al-Hadi, Mohammed A. Esmail, Jamil Al-Jurban, Abdulsamad Alkawri, Arif Shamsan, Payton Gomein, Jeanne N. Samake, Adel Aljasari, Abdullah A. Awash, Samira M. Al Eryani, Tamar E. Carter

## Abstract

*Anopheles stephensi* is an invasive malaria vector in Africa. To determine the status of the mosquito in Yemen, *An. stephensi* vector surveillance and molecular confirmation was conducted in Al Hudaydah Governorate in 2021 and 2022. Mosquito larvae were collected in suspected man-made breeding habitats in Ah Dahi and Zabid city in 2021 and 2022, respectively. Mosquitoes morphologically identified as *An. stephensi* underwent molecular confirmation through PCR assays, sequencing, and phylogenetic analysis of the cytochrome oxidase subunit I (COI) gene and internal transcribed spacer 2 locus (ITS2). Analysis confirmed *An. stephensi* identification for the majority of samples (39/41), with two COI haplotypes detected: one newly reported haplotype and one haplotype common to Northeast Ethiopia and Somaliland. No clustering with *An. stephensi* from the Arabian Peninsula was observed. These findings provide preliminary insight into the diversity of *An. stephens*i in Yemen and the connection between *An. stephensi* in Yemen and East Africa.

## Background

Malaria remains one of the most significant threats to global health, with about 247 million cases reported in 2021 (WHO 2022). On-going challenges including antimalarial (1) and insecticide resistance (2) remain a consistent threat to progress. In recent years, a new challenge has emerged with the invasive malaria vector *Anopheles stephensi* in East Africa. *Anopheles stephensi* was believed to be restricted to South Asia, the Middle East, and portions of the Arabian Peninsula (3). However, since the first detection in Africa specifically in Djibouti in 2012 (4) and in eastern Ethiopia in 2016 (Carter et al. 2018)(5), *An. stephensi* has been found in Somalia (6), Sudan (7), Nigeria (8), Eritrea (8), Kenya (9), and Ghana (10). With the mounting evidence of its resistance to multiple classes of insecticides (11, 12) and its association with recent malaria outbreaks (13), concerns about its status and spread in the Mediterranean region are heightened.

In the Arabian Peninsula, the distribution of *An. stephensi* is unclear. Previous predictive models and field investigations indicate native populations exist in the northeastern coastal region along the Persian Gulf and inland in countries including Saudi Arabia (Sinka et al 2011)(3). The status of *An. stephensi* is of particular importance in Yemen, where a notable increase in malaria cases has been reported in Aden City starting in 2017 (14). In Yemen, the primary vector is *Anopheles arabiensis*. The first report of *An. stephensi* in Yemen occurred in Aden City, southern Yemen in 2021 (8) followed by the molecular confirmation in 2023 (Allan et al., 2023). However, recent retrieval of an unpublished entomological survey report included *An. stephensi* in 2000, in sites within one district (Al Zuhra) in Al-Hudaydah Governorate, western Yemen (WHO Yemen, unpublished report, 2000). No record of *An. stephensi* was documented before 2000 or in following entomological surveys until the recent detection in Aden City in 2021. To date, little is known about the distribution and characteristics of this vector in western governorates where the highest burden of malaria in the country is found (14). *Anopheles stephensi* was reported for the first time in Ad Dahi and Zabid districts within the Al Hudaydah Governorate in December 2021 and March 2022 respectively. Since then *An. stephensi* has been found in multiple suburban areas in the Tehama coastal plain region (15). However, this study focused on the molecular confirmation of *An. stephensi* found during vector surveillance in Ad Dahi and Zabid districts in the Al Hudaydah Governorate.

## Methods

### Site Descriptions

Ad Dahi is located north of Hodeida city and is semiurban (15° 12’ 55” N / 43° 4’ 13” E) (Figure 1). The population size of Ad Dahi town is about 21,587 and the climate is tropical with a high grade of temperature in the summer and moderate in the winter. Zabid is in one of the southernmost districts of Al Hudaydah governorate located in the Tehama coastal plain near the Red Sea (14°12’02.6”N, 43°19’06.6”E) in a tropical climate (Figure 1). Zabid town is also semiurban with a population size of 34,686.

**Figure 1.**
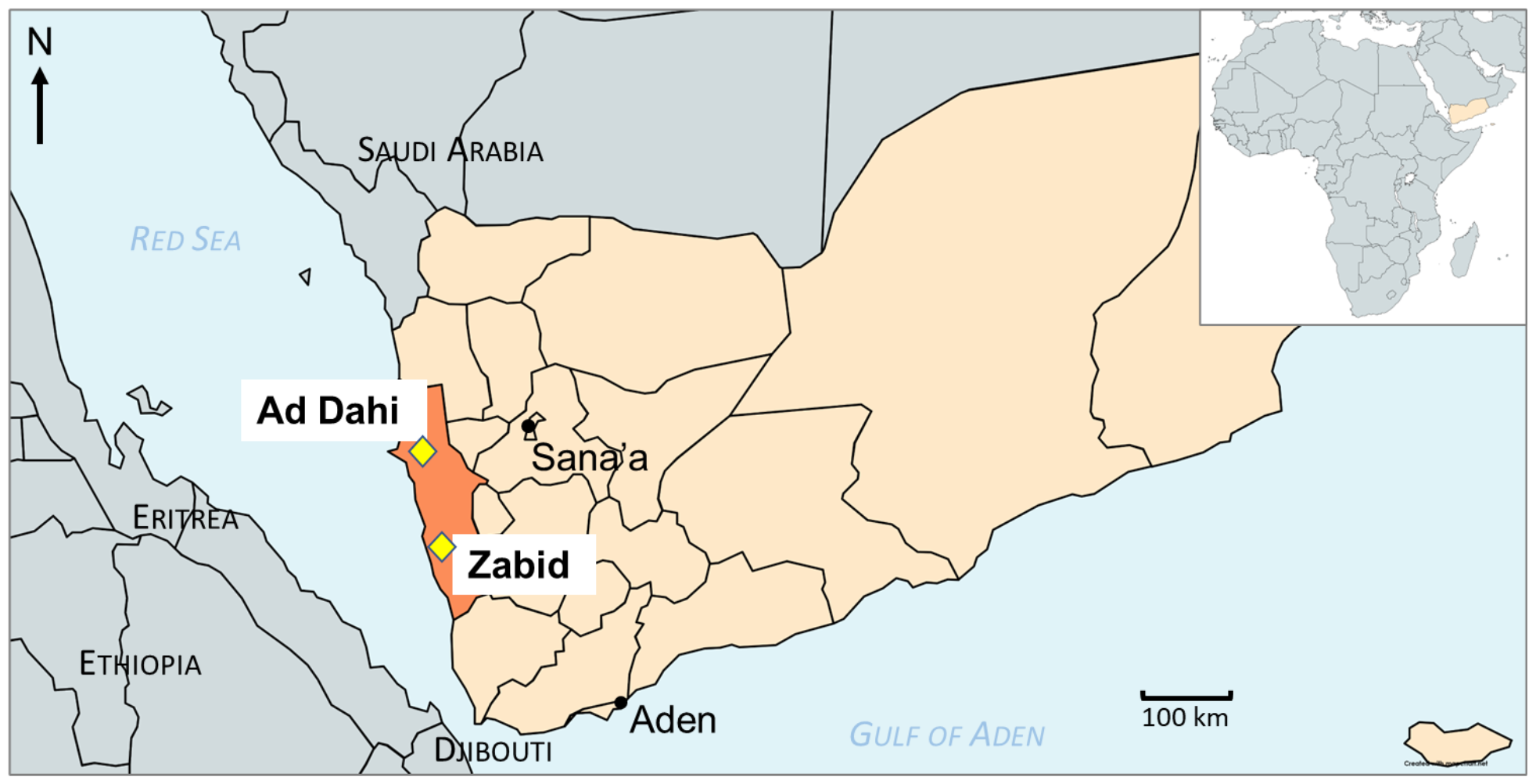
Map of Yemen with locations where *An. stephensi* was detected in 2021-2022, marked with yellow diamond.

### Collections

Mosquito larvae were collected using the dipping method in one day in December 2021 in Al Dahi and in March 2022 in Zabid district as part of *Aedes* and *Anopheles* surveillance, respectively. The Al Dahi survey was a response to a recent dengue virus outbreak in that region. Potential breeding containers surveyed included open concrete ponds, and cement water tanks that are used for multiple purposes, such as block factories, washing in mosques, and car washing places, etc. Mosquito larvae were reared in field insectary to adults. Morphological identification was determined using a recently updated key (16). The specimens morphologically identified as *An. stephensi* were preserved with silica gel and a subsample of specimens was sent to Baylor University for molecular analysis.

### Molecular and sequence analysis

Two loci were selected for PCR-based species identifications: cytochrome oxidase subunit 1 (*COI)* and internal transcribed spacer 2 (*ITS2)*. Analysis of *COI and ITS2 have* been used for *An. stephensi* species confirmation in an invasion context in previous studies (6, 17). First, a stephensi-specific PCR endpoint assay was used to identify *An. stephensi* based on amplification (presence/absence) of a portion of the ITS2 locus. The primer sequences for PCR in the *ITS2* endpoint assay were 5.8SB (5′-ATCACTCGGCTCGTGGATCG-3′) and 28SC (5′-GTCTCGCGACTGCAACTG-3′) (18). Then two additional PCR protocols for the *ITS2* locus and the *COI* gene were implemented to generate products for sequencing. PCR was conducted as detailed in Carter et al. (5). The primer sequences for the *ITS2* PCR for sequencing were 5.8SB (5′-ATCACTCGGCTCGTGGATCG-3′) and 28SB (5′-ATGCTTAAATTTAGGGGGTAGTC-3′) (18). The primer sequences for *COI* PCR were LCO1490F (5′-GGTCAACAAATCATAAAGATATTGG-3′) and HCO2198R (5′-TAAACTTCAGGGTGACCAAAAAATCA-3′) (19). *COI* and *ITS2* PCR products were sequenced using Sanger sequencing technology. Sequences were then trimmed using CodonCode Aligner (CodonCode Corporation, Centerville, MA) and submitted to the National Center for Biotechnology Information’s (NCBI) Basic Local Alignment Search Tool (BLAST) to confirm successful amplification. The COI sequences were further aligned in CodonCode with previously published sequences retrieved from Genbank, and phylogenetic analysis was conducted using maximum-likelihood method with RAxML (20). The final trees were annotated using Figtree (21).

## Results and Discussion

The majority of the *Anopheles* larvae were collected from cement water tanks found outside the homes. Forty-one mosquitoes were reared to adult stage, identified morphologically as *An. stephensi*, and analyzed molecularly (7 from Ad Dahi and 34 from Zabid City). Of the 41 samples, all 7 Ad Dahi and 32 Zabid City specimens were confirmed to be *An. stephensi* with ITS2 end-point assay and by COI targeted sequence analysis. Of the two non-*An. stephensi*, one was determined to be *Anopheles culicifacies* and another *Aedes aegypti* based on *ITS2* and *COI* sequence analysis. Phylogenetic analysis confirmed the species identification of the *An. stephensi* specimens (Figure 2 and 3).

**Figure 2.**
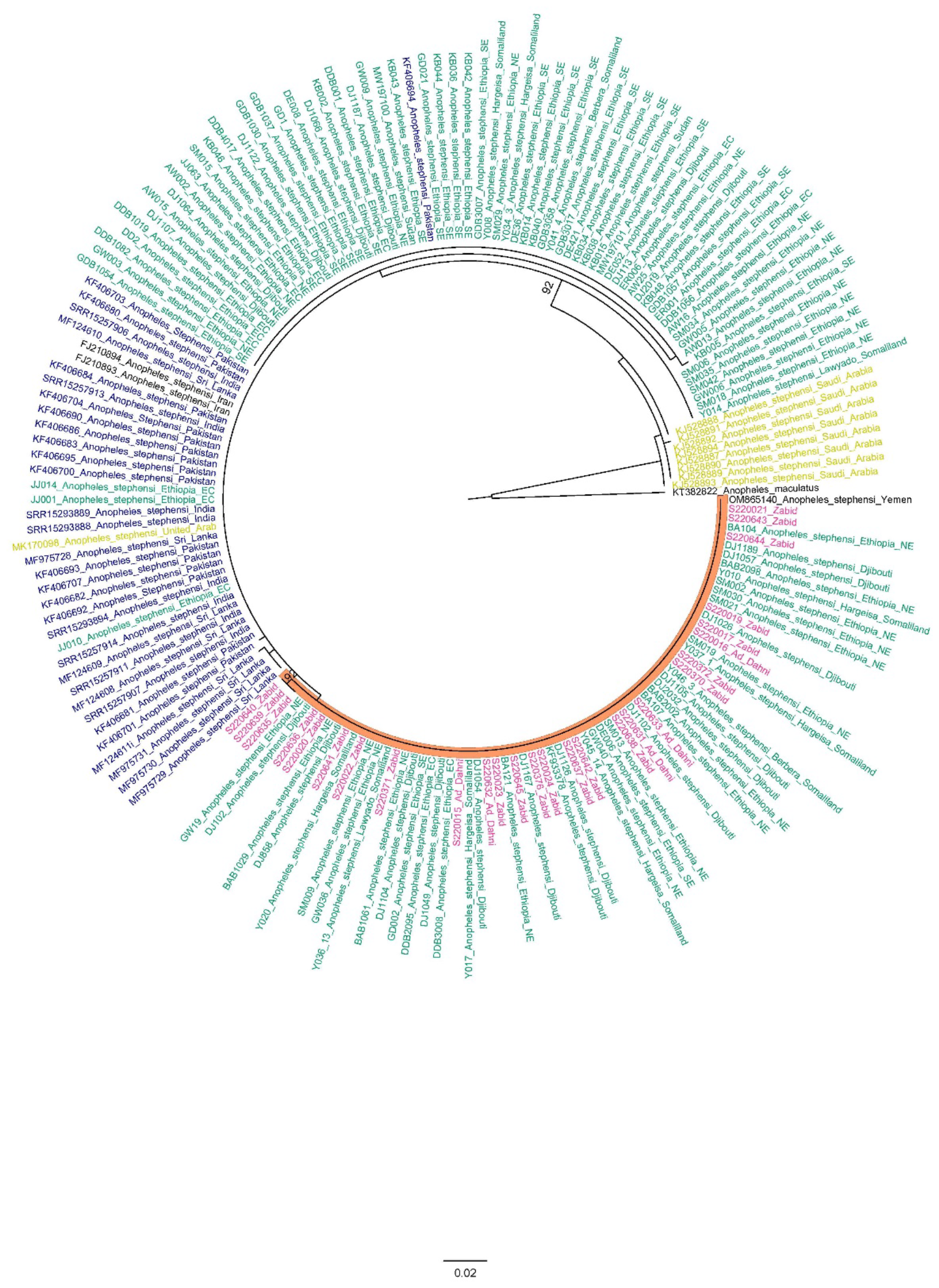
Phylogenetic analysis of *An. stephensi* COI sequences in Yemen using maximum likelihood approach. Yemen sequences are in pink. South Asian are blue, Horn of Africa are green, and Arabian Peninsula are yellow-green. Orange highlight indicates branch containing Yemen and Horn of Africa specimens only.

**Figure 3.**
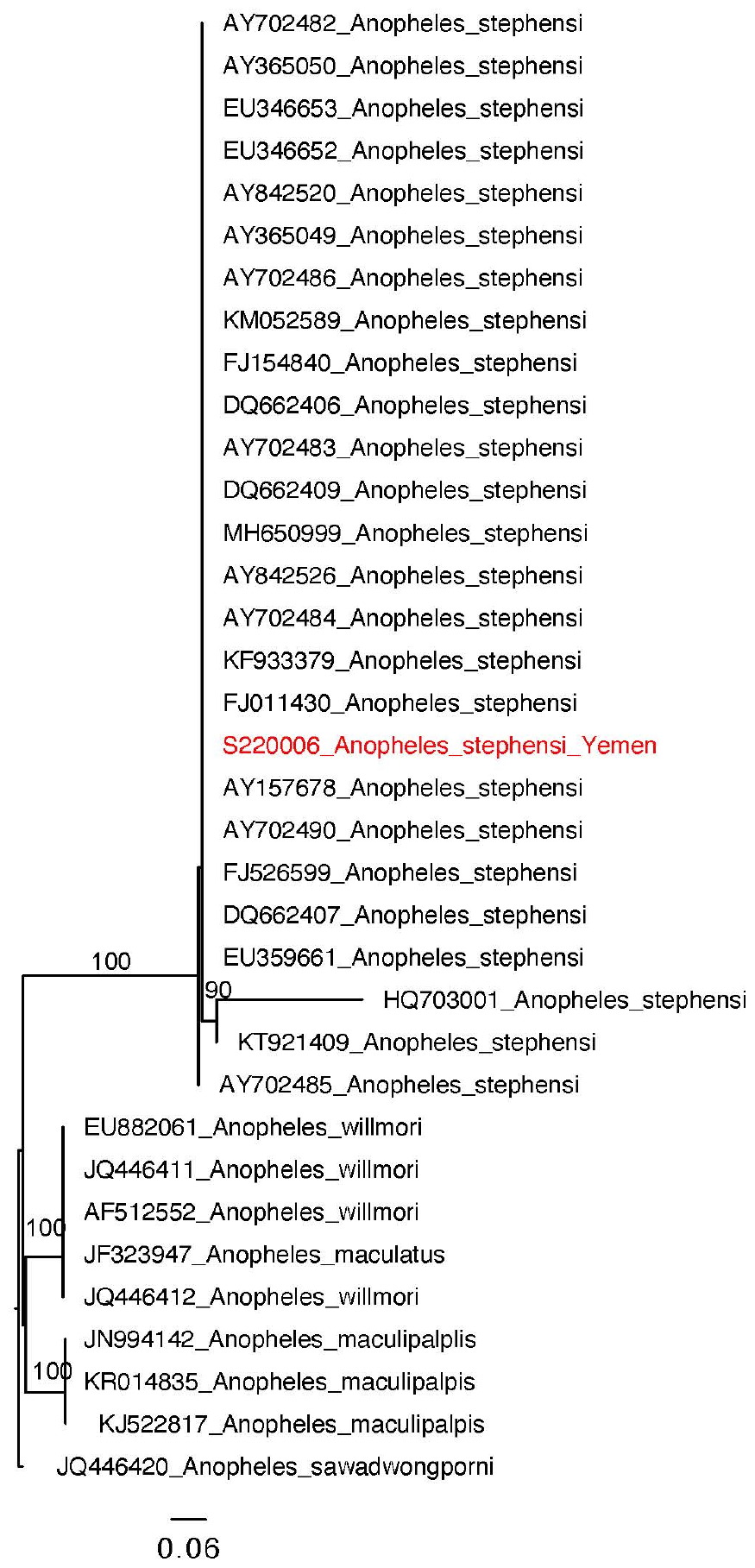
Phylogenetic analysis of *An. stephensi* ITS2 sequence from Yemen using maximum likelihood approach. Only one sequence is included representative of the single haplotype observed in Yemen *An. stephensi*. Yemen sequence is highlighted in red. Only bootstrap value above 70 are included.

ITS2 sequences were all identical for *An. stephensi* specimens and BLAST revealed a 100% sequence identification for *An. stephensi* in GenBank. Sequence analysis of COI revealed two haplotypes. One haplotype (n= 35) was previously reported throughout the Horn of Africa (COIHap 3)(22) with the highest frequency in northeastern Ethiopia, Somaliland, and Djibouti. This haplotype was also detected in a recent report on the detection of *An. stephensi* in Aden, Yemen (23). The second haplotype (n=4), differing by a single nucleotide from COIHap3, has not previously been reported.

These findings serve as the first molecular confirmation of *An. stephensi* in the Al Hudaydah Governorate and demonstrate the successful utilization of ongoing vector surveillance activities for the initial detection of *An. stephensi*. The most productive breeding habitats were cement water storage tanks as is commonly observed in other invasive settings in eastern Ethiopia (5, 24), and Aden, (23). The detection of a common Horn of Africa *COI* haplotype raises questions about the relationship between the invasive *An. stephensi* in northern Horn of Africa and Yemen. It is possible that they both share a common origin or that movement of *An. stephensi* has occurred or is occurring between the two regions. A hypothesis can be generated from these findings about the relative timing of Yemen introduction relative to the HoA based on the following observations. 1) Fewer haplotypes were observed in Al Hudaydah district and in Aden compared to Northern Ethiopia, Somalia, and Djibouti, which may be indicative of a more recent introduction in Yemen. 2) Hap3 has a limited geographic range compared to the HoA-wide Hap2, which suggests this haplotype could be associated with a later introduction in HoA and potentially Yemen. Furthermore, as with the Horn of Africa specimens and the study in Aden, no Saudi Arabian haplotypes were detected. Moreover, unlike the Horn of Africa, no South Asia haplotypes were detected. Thus, further genomic analysis and extensive *An. stephensi* sampling in Yemen *An. stephensi*, Saudi Arabia, and other parts of the Arabian Peninsula are needed to evaluate the hypothesis of a more recent introduction in Yemen relative to the HoA.

## Conclusion

These findings provide the first molecular confirmation and genetic diversity of *An. stephensi* populations in Al Hudaydah Governorate, Yemen. The findings also provide support for the need of additional long-term entomological and epidemiological surveillance of the impact of this invasive malaria vector on malaria transmission in the region. With the detection of *An. stephensi* in Yemen, questions linger related to its relationship to the HoA *An. stephensi*. Further genomic analysis should be performed to examine the hypothesis proposed from the results of this study.

## Availability of data and materials

The sequences generated in this study are available through NCBI Genbank Accession numbers pending.

## Abbreviations

HoA: Horn of Africa
WHO: World Health Organization
PCR: polymerase chain reaction
ITS2: internal transcribed spacer 2
COI: cytochrome c oxidase I
NCBI: National Center for Biotechnology Information
BLAST: basic local alignment search tool
SNPs: single nucleotide polymorphisms

## Acknowledgments

The authors would like to thank Dr. Ghasem Zamani (Regional Advisor WHO/EMRO-MVC) for his contribution to the conceptualization of the project, technical support for implementation of surveillance and review of the manuscript. We would also like to thank Yamaan Foundation for Health and Social Development for their support of this work.

## Contributions

TEC, MA, and SAE contributed to the conception and design of the project. MA, MAH, MAE, JAJ, AS organized and led the collection of specimens. TEC, PA, JS generated the data. TEC and PA analyzed the data. TEC wrote the first draft of the manuscript. All authors read and approved the final manuscript.

## Biographical Sketch

Tamar Carter is an Assistant Professor of Tropical Disease Biology at Baylor University in Waco, Texas, USA. Her research interests include evolutionary genomics of invasive mosquito vectors and host-parasite coevolution.

Competing Interests:

The authors have no competing interests.

## Consent for Publication

This manuscript is published with permission of all authors.

## Participation of Human Subjects

There were no human subjects involved in this Synopsis.

### Disclaimer

The authors alone are responsible for the views expressed in this article and they do not necessarily represent the views, decisions, or policies of the institutions with which they are affiliated.

